# Surface Transmon Resonance (STR): a handheld nanogap biosensor for real-time, label-free molecular binding kinetics

**DOI:** 10.64898/2025.12.29.696853

**Authors:** Bryan K Chantigian, Sang-Hyun Oh

## Abstract

Despite the exponential growth of microelectronics driven by Moore’s law over half a century, biosensing has remained largely dominated by optical methods such as fluorescence assays and surface plasmon resonance (SPR). Electronic biosensors, particularly field-effect transistor (FET)-based platforms, have historically faced challenges related to Debye screening and the lack of compact, high-frequency instrumentation necessary for label-free, high-sensitivity measurements in conductive solutions. Here, we present a Surface Transmon Resonance (STR) biosensor; a nanogap-enabled electronic platform that exploits a handheld vector network analyzer (nanoVNA) for real-time, phase-resolved detection of protein–protein interactions. Operating at radio frequencies in the hundreds of MHz range, STR decreases Debye screening limitations and provides an SPR-like readout with significantly reduced cost and footprint compared to benchtop optical systems. By characterizing specific bovine serum albumin (BSA) and anti-BSA binding kinetics, we demonstrate that STR biosensors can quantitatively assess molecular affinity and interaction dynamics, capabilities traditionally reserved for optical gold-standard techniques like SPR. This work positions STR as a compact, scalable, and cost-effective electronic alternative to conventional optical biosensors.

## Introduction

Optical biosensors, particularly surface plasmon resonance (SPR) and fluorescence-based methods, have dominated biomolecular interaction studies [1–7]. However, their reliance on bulky optics, expensive instrumentation, and the need for precise alignment presents limitations for cost-effective, portable point-of-care (POC) diagnostics [8]. Electronic biosensors, particularly field-effect transistor (FET)-based platforms, have been considered as an alternative, offering label-free sensing and scalable chip integration [9–14]. Even so, their performance is hindered by Debye screening in physiological solutions and the weaker confinement of RF fields compared to the highly localized evanescent fields in SPR, resulting in reduced surface sensitivity [15–17].

**Table 1:** Comparison of typical sensor architectures.

|  | SPR<br>(gold standard) | STR<br>(this work) | ISFET | EIS |
| --- | --- | --- | --- | --- |
| Debye Screening Limited | Optical sensor, not affected | No (High-frequency operation, >100 MHz) | Yes | Somewhat (0 Hz - 1 MHz) |
| Real-Time Kinetics | Yes | Yes (phase-resolved) | Limited (drift-sensitive, poor S/N) | Limited (slow freq sweep) |
| Portability | No | Yes | Yes | Yes |
| Cost | \$100,00 - \$500,000 | \$300 | \$500 - \$2,000 | \$5,000 - \$25,000 |

In an attempt to overcome the limitations of existing biosensing architectures, we introduce Surface Transmon Resonance (STR), a low-cost, label-free electronic biosensing platform based on high-frequency transmission-line resonance (Fig. 1). The STR architecture features a nanoscale gap between coplanar transmission lines, forming a resonant tank circuit with surface-confined electric fields highly sensitive to local permittivity changes induced by biomolecular interactions [18–22]. Unlike conventional RLC tank circuits or microelectrode-based designs [23], the STR architecture uniquely combines nanogap field enhancement, high-frequency RF operation, and real-time, phase-sensitive readout. The architecture for STR is inspired from transmon qubits, as these use an electronic resonance to interact with an object under study within a gap region in the transmission lines. This interaction affects the device’s resonance frequency. In the case of transmon qubits, the objects under study are Josephson junctions whereas our architecture utilizes biomolecules as the object under study. Operating at hundreds of MHz, STR mitigates Debye screening effects and enables label-free biomolecular detection in physiologically relevant solutions [15, 24–26]. The system integrates seamlessly with a handheld vector network analyzer (nanoVNA), offering a compact, scalable solution for quantitative molecular interaction analysis.

**Figure 1:**
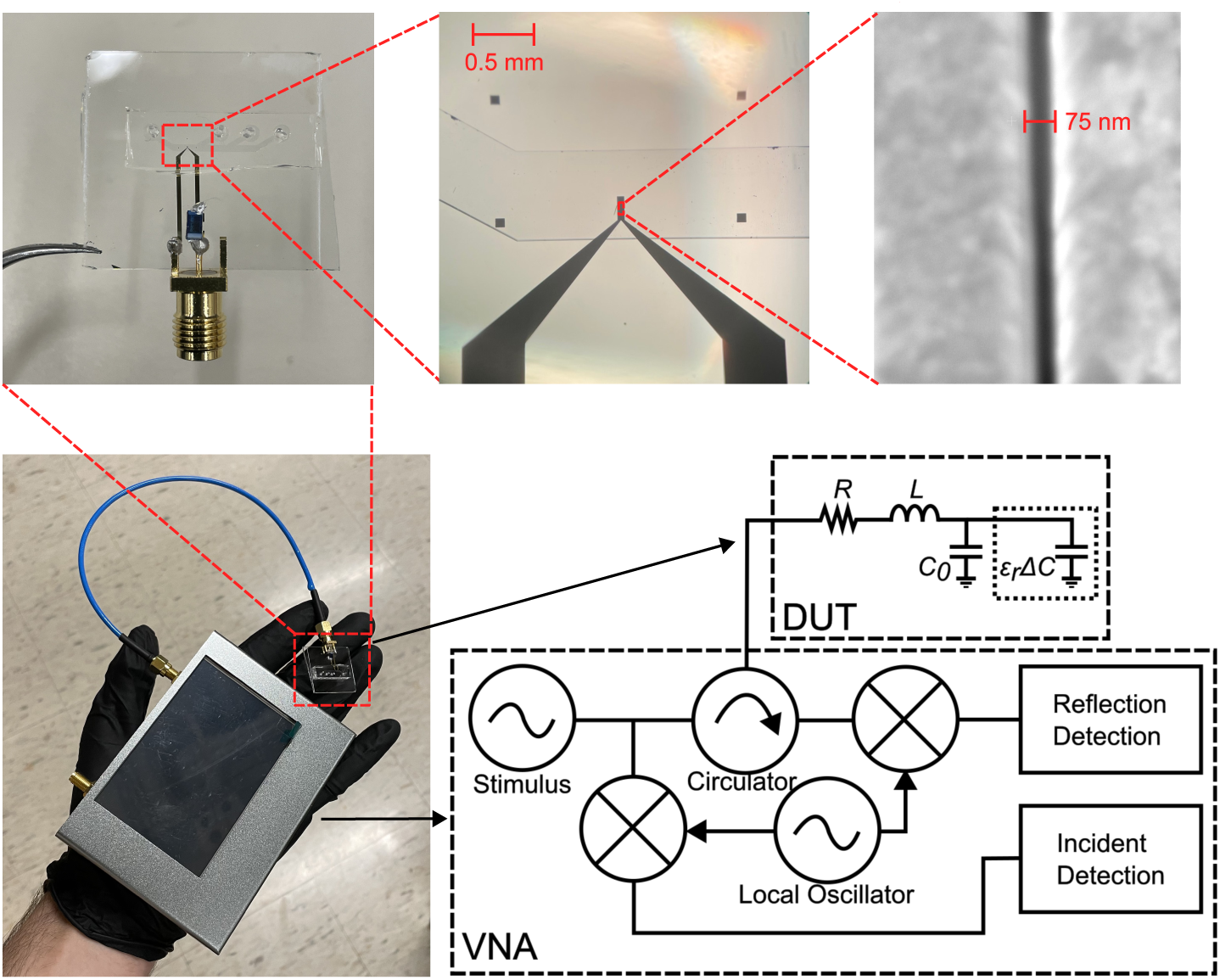
Sensing system of the VNA connected to the nanogap sensor (bottom left), a magnified image of the nanogap sensor (top left), microscope image of sensing region at 5x magnification (top middle), an SEM image of the nanogap region (top right), and the circuit schematic (bottom right) of the system including a generic VNA circuit diagram and the nanogap sensor (DUT).

By quantifying bovine serum albumin (BSA) and anti-BSA binding kinetics - a benchmark traditionally measured using gold-standard SPR - we demonstrate that STR provides a rapid, scalable, and cost-effective electronic alternative to optical biosensors. This protein-ligand system is used to show proof of concept of this new architecture and to provide groundwork for more clinically relevant samples in the future. With its simplified electronic readout, compact form factor, and low-cost implementation, STR overcomes the limitations of bulky and expensive optical systems, making it a promising platform for real-time, label-free biomolecular analysis and POC diagnostics. This work presents a new class of portable electronic biosensors that combine nanofabrication, high-frequency RF circuit design, and label-free surface-based detection. This architecture paves the way for scalable, real-time biomolecular diagnostics outside traditional laboratory settings by decreasing system cost and complexity while maintaining high sensitivity in a compact footprint.

## Results

### High-frequency resonance enables analysis of molecular interactions not limited by the Debye length

The sensor developed in this work is designed as a resonant circuit. The sensing portion consist of small open circuit transmission lines separated by a nanogap. This sensing region is then connected in series to an inductor, forming a series RLC circuit shown in Fig. 1. The resonant architecture is particularly advantageous, as it restricts the measurement to a narrow frequency range, providing clear, well-defined signal changes in response to biomolecular interactions, similar to SPR. Unlike sensor systems such as electrochemical impedance spectroscopy (EIS), which analyzes a broad frequency spectrum - offering rich and robust information but complicating data interpretation - our STR approach leverages resonance for simplified, high-contrast sensing [27–31].

The sensor’s resonance frequency can be determined by 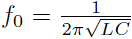. From here, it is obvious that changes in either the inductance or capacitance cause the resonance frequency to shift.

The total capacitance of the sensor system is determined by parasitic capacitance, *C*_0_, and the change in capacitance due to localized permittivity changes, *ɛ_r_*Δ*C*, and results in *C* = *C*_0_ + *ɛ_r_*Δ*C*, where the values are simply added.

Typically, conductive solutions will form an electric double layer (EDL) causing a large increase in capacitance while simultaneously limiting the penetration of the sensor’s electric field, affecting the effective permittivity measured [32, 33]. In this sensor’s design, this EDL would form between the gold electrodes shown in Fig. 2. EDL screening can be a significant hurdle in biological sensing, as biological samples must be maintained in buffered ionic solutions to ensure stable pH and physiological conditions. The advantage of this sensor’s architecture is that the frequency of operation is in the hundreds of MHz. At this frequency, the ions in solution have difficulty responding to the rapidly changing field, weakening the formation of an EDL and thus the screening effect by the ions in solution is decreased [15, 24, 26, 34–36]. Simulations from Pittino et al. [24], determine the frequencies in which the capacitance introduced from the EDL is decreased. When this capacitance values is lowered, it indicates that the EDL is not as prevalent. This frequency value was found to scale roughly linearly with the salt concentration of the solution. For example, they state that the EDL capacitance cutoff frequencies for 5 mM and 500 mM NaCl concentrations are around 15 MHz and 1.5 GHz, respectively. Extrapolating from these values, for the buffer used in this study, 0.1x PBS which has a NaCl concentration of around 13.7 mM, the expected EDL capacitance cutoff frequency would be approximately 40 MHz.

**Figure 2:**
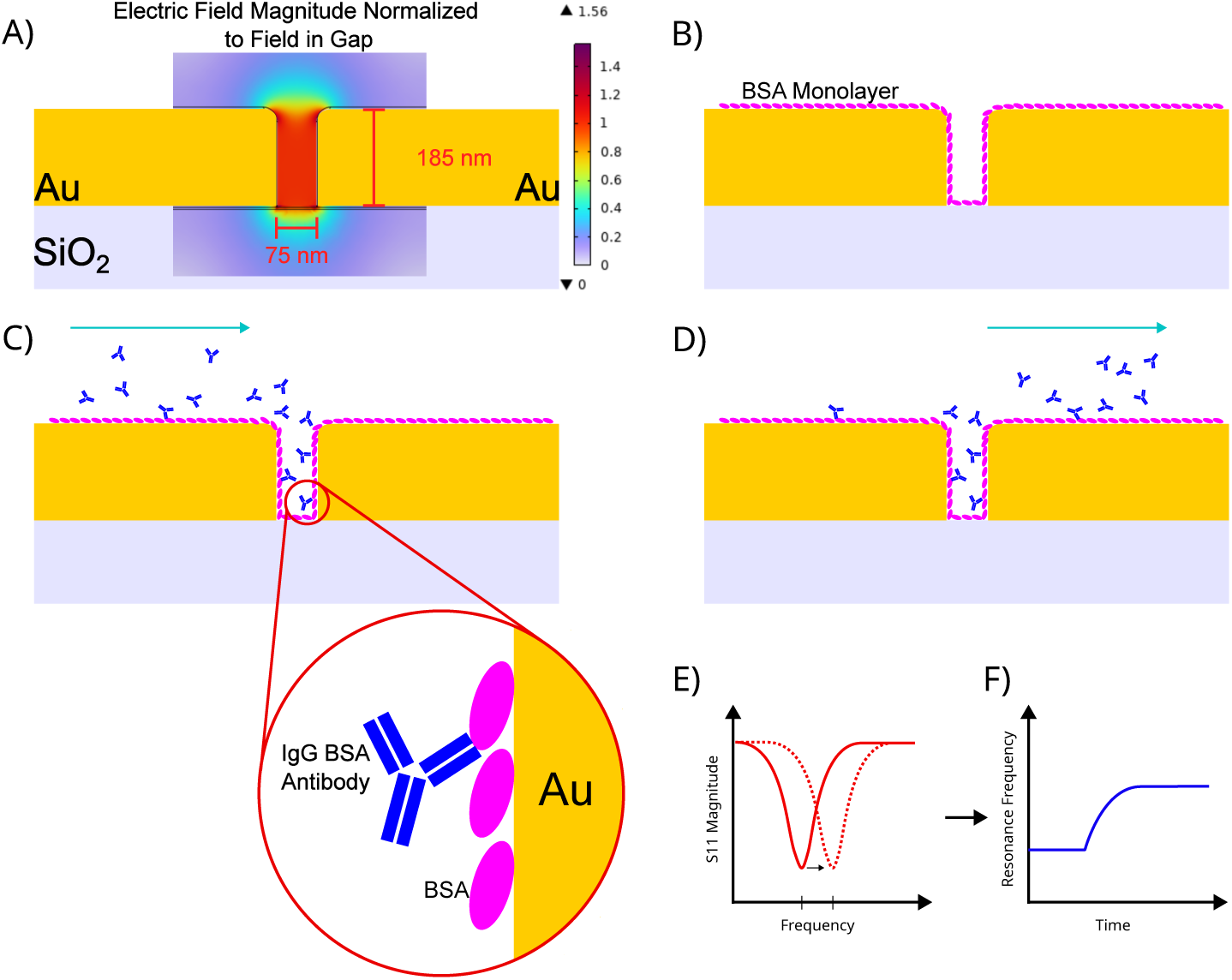
STR sensing process at the nanoscale. The light blue arrow indicates the flow of the solution. A) shows the sensing region before any binding occurs; Overlaid COMSOL simulation (at 150 MHz) shows electric field confinement in the nanogap region. B) Injection of bovine serum albumin (BSA), which adsorbs to gold surfaces inside the nanogap. C) Introduction of anti-BSA antibodies, which selectively bind to surface-adsorbed BSA. D) Buffer is reintroduced to wash out unbound antibodies, allowing dissociation dynamics to be monitored. E) Accumulation of molecules inside the nanogap perturbs local permittivity, shifting the resonant frequency. F) Real-time tracking of resonance frequency shift captures kinetic binding curves over the measurement period.

### Nanogap-enabled surface sensitivity in high-frequency biosensing

The sensor uses the change in the nanogap’s capacitance in order to determine the presence of proteins. Since the size of the gap region is on the same order of magnitude as many common proteins, these biomolecules take up a considerable volume within the gap, thus changing the sensor’s resonance frequency [37]. An illustration of this can be seen in Fig. 2.

It has been highlighted that this sensor primarily measures permittivity but it is also able to measure a solution’s conductance. Measurements of capacitance - reflecting the relative permittivity - allow for the binding and unbinding of various biomolecules to be observed, whereas conductance measurements allow ion concentrations to be determined. An analytical equation for determining this is given in the Supplementary Material. Typically, during the tests highlighted in this paper, the solutions used were of similar conductivity in order to more accurately determine protein binding kinetics. When conductivity changes were observed, they were typically calibrated out as they occurred almost instantaneously and the solutions’ ion concentrations were already known.

### Phase-resolved detection for robust biomolecular sensing

The sensors had an *S*_11_ Q-factor between 50-100 when they were immersed in deionized water, as shown in Figure 3. Although, when using biological buffers, the *S*_11_ Q-factor can become degraded due to the increased conductance of the solution causing mismatch between the VNA’s impedance and the load impedance. This can make it more difficult to determine properties correlated with permittivity, but phase-based measurements were found to help elucidate the resonance frequency shift. The signal to noise ratio (SNR) of the *S*_11_ phase-based measurements were found to be higher than the SNR of the *S*_11_ magnitude-based resonance shift measurements. Since these phase-based measurements are more robust, they are primarily utilized in this work to verify and determine biomolecular binding kinetics.

**Figure 3:**
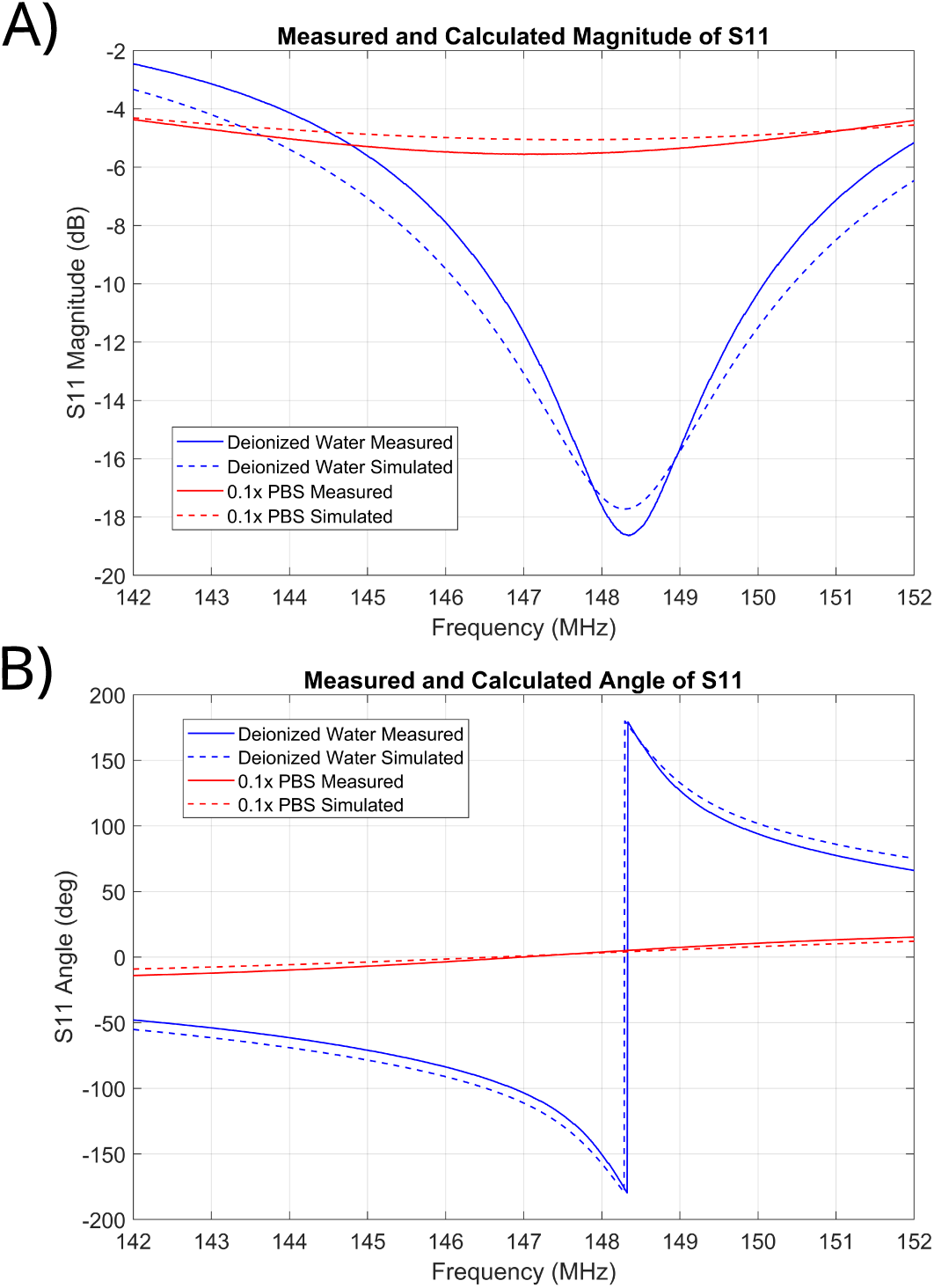
The Q factor of *S*_11_’s magnitude. (A) is decreased for higher conductance samples since the real part of the impedance is increased. The minimum magnitude is shifted to a lower frequency due to the increase in conductance. The slope of *S*_11_’s angle (B) is decreased when the conductance of the sample increases, increasing the real part of the impedance. The location where the *S*_11_ angle passes zero is shifted since the conductance increases. The sudden jump from -180 to 180 degrees for the deionized water is due to the phase being wrapped. The phase passes through 180 degrees since the real impedance is less than the characteristic impedance of the VNA.

In addition to picking the peak magnitude and phase zero-crossing, calculating the centroid of the resonance dip will also allow for enhanced determination of the resonance frequency. From the test conducted, the accuracy of the centroiding method appears to be on-par with the phase zero-crossing methodology. Phase-based calculations were utilized as they were less computationally intensive and do not require as much data. This allows for phased-based calculations to collect a narrower band of frequencies which, then, allows for the VNA’s scan speed or resolution to be increased. For computation of the resonance frequency, a simple first order linear model is fitted to the phase returned around zero, and the equation is then used to determine a more precise zero crossing point. This allows for higher precision in determining the resonance frequency, compared to choosing a data point near zero.

### Demonstration of biomolecular kinetic binding measurements

In order to test the nanogap sensor, a procedure, shown in Fig. 2, was created. To get a baseline reading, a buffer solution without any biomolecules was injected into the biosensor’s microfluidic channel. After, bovine serum albumin (BSA) was injected so that the BSA molecules would bind to the sensor. Once the surface was saturated with BSA, the channel was flushed with the buffer and BSA antibodies were then injected so binding may be seen. When equilibrium was achieved, the buffer was then injected into the channel in order to detect unbinding.

The shifts in resonance frequency, due to local permittivity change from proteins, can be seen in Fig. 4. These curves were used to observe protein binding kinetics and calculate association and dissociation rate constants. The resonance frequency of a nanogap sample was measured at the same time as a sensor that had a 10 *µ*m gap, as this would verify that the changes in frequency were indeed due to surface interactions and not to properties of the bulk. Surface interactions are not measurable in the microgap sensor primarily because the total effective volume of the proteins in the gap is negligible compared to the total volume of the 10 *µ*m gap, whereas the nanogap’s volume is on the order of the effective volume of the proteins so surface sensitivity is enhanced. In the case of the BSA binding, the layer formed by the BSA on the electrodes is approximately 1-4 nm [38, 39]. This would take up around 3-20% of the nanogap’s volume, whereas it would take up only around 0.02-0.08% of the microgap’s volume. When there is only the buffer and no biomolecules present, the response of the sensor stays reasonably flat. Once the BSA solution is injected into the microfluidic channel, the response of the 10 *µ*m sensor jumps, while the nanogap sensor rises over time due to the BSA binding to the surface of the nanogap. After the BSA blocking step and flushing the channel with buffer solution, BSA antibodies were injected into the channel. Both sensors have similar responses to when the BSA solution was injected, the microgap’s resonance jumps up again due to the overall permittivity change while the nanogap’s resonance rises more slowly due to the binding of the antibodies to the BSA adsorbed to the surface.

**Figure 4:**
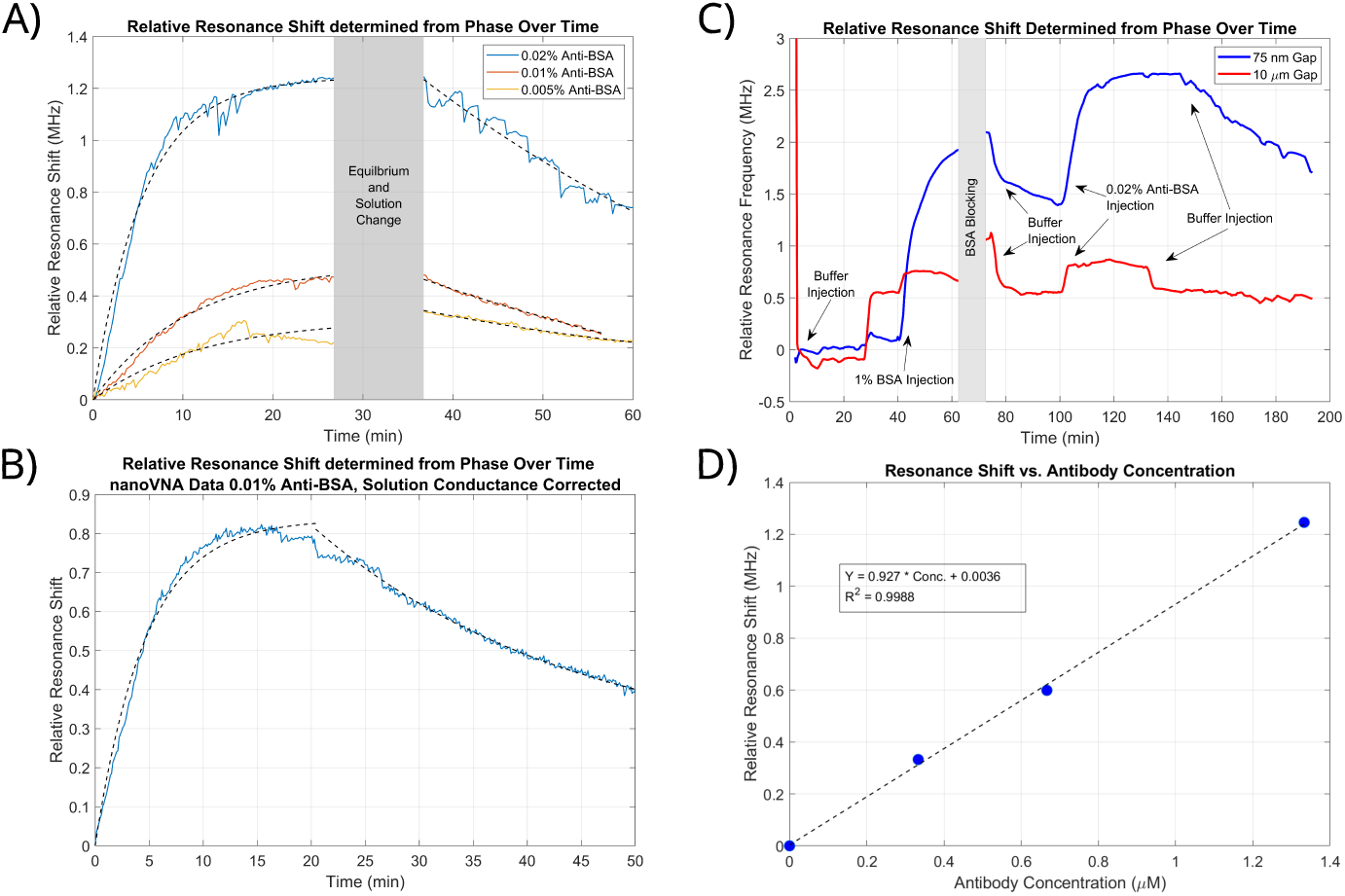
Resonance shift over time of. A) Anti-BSA with concentrations of 0.02%, 0.01%, and 0.005% binding to the 75 nm gap sensor, B) Anti-BSA binding measured by nanoVNA, and C) binding measured at 75 nm gap sensor and 10 *µ*m gap sensor. D) Calibration curve for the STR sensor showing the relative resonance shift over antibody concentration. The resonance frequency for each of these plots was determined using the measured *S*_11_ phase.

The process of BSA blocking and injecting BSA antibodies was repeated on different sensors at different concentrations. This allowed for determination of the binding kinetic parameters of the tested BSA antibody using the equations listed below [40]:

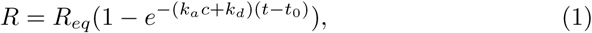

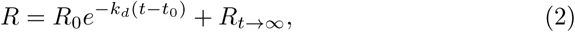

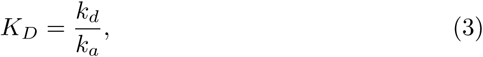

where *R* is the response of the sensor, *R_eq_* is the response at equilibrium, *k_a_* is the association constant with units *M^−^*^1^*s^−^*^1^, *c* is the concentration of the antibody in *M*, *k_d_* is the dissociation constant with units of *s^−^*^1^, *t* is the time in seconds, and *t*_0_ is the time offset when binding begins. In equation (2) *R*_0_ is the amplitude of decay and *R_t→∞_* is the offset response due to any non-specific binding between the antibody and the sensor. Finally, in equation (3), *K_D_* is the equilibrium dissociation constant. Binding and unbinding curves of various different anti-BSA concentrations is shown in panel A of Fig. 4. These curves were used to determine the equilibrium dissociation constants for the specific antibody being used. The equilibrium dissociation constants for the 0.02%, 0.01%, and 0.005% concentrations were 143 nM, 135 nM, and 145 nM, respectively. The equilibrium dissociation constant determined from our SPR system was around 130 nM, showing good agreement between the STR and SPR systems. For further information, the SPR curve and analysis of SPR binding kinetics are discussed in the Supplementary Material of this paper.

Mass transport limitations (MTL) can cause error in determining kinetic binding parameters. Instances in which biomolecules are not able to quickly move away from the sensing surface, MTL, can cause the biomolecular concentration near the surface to decrease in the association phase and increase in the dissociation phase. This causes the kinetics to appear slower than they actually are [41]. To further ensure the protein kinetics measured by the STR sensor are not limited by mass transport, the Damkohler number can be used. The Damkholer number compares the reaction rate to the mass transport rate which, in this case, is driven by the biomolecular binding and the fluid flow. When this number is much less than 1, the reaction is said to be reaction rate limited and not mass transport limited. In this case the number is determined from the association rate, *k_on_*, the maximum concentration of biomolecules tested, *C*_0_, the fluid velocity at the height of the sensor, *v*, and the width of the electrodes, *w*. This gives the equation, 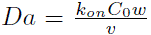. Assuming laminar flow within the channel, the flow velocity at the height of the sensor is calculated to be approximately 80 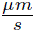 at a pump rate of 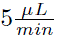. The association coefficient was determined from SPR to be approximately 3773 *M^−^*^1^*s^−^*^1^, and the maximum concentration tested was 1.33 *µM*. With all of these values, in addition to a total sensor electrode width of 60 *µm*, the Damkohler number is calculated to be approximately 0.004. This value, in conjunction with the agreement of equilibrium dissociation constants between STR and SPR, strongly indicates that the protein binding kinetics are not mass transport limited.

## Discussion

The BSA/anti-BSA demonstration validates STR as a general-purpose biosensing architecture capable of quantifying specific molecular interactions in real time. Its SPR-like kinetic curves and phase-resolved signal clarity position it as a viable alternative for applications such as drug screening, biomarker quantification, and field-ready diagnostics. Crucially, this performance is achieved with a palm-sized device, dramatically reducing cost and complexity compared to conventional benchtop SPR systems. The nanoVNA, initially released in 2019, allows for the significant decrease in price, as typical VNAs can cost thousands of US dollars, and allows for the miniaturization of the system since it is a handheld VNA [42–44]. Although the nanoVNA provides a significant advantage to making the system affordable and portable, it is certainly possible to measure these binding events with other VNAs, as the key for observing this is the nanogap sensor itself. The operation of the nanogap sensor was verified using a compact, USB VNA from Keysight (Model # P9372A). Resulting curves are shown in panel A and C of Fig. 4. The nanoVNA was tested as well and showed a similar binding curve, panel B of Fig. 4, and the measurement resulted in an equilibrium dissociation constant close to other values measured at 158 nM.

This sensor system’s linear sensitivity can be determined from the expression for the resonance frequency and total capacitance. When plugging those equations into each other and then taking it’s Taylor expansion, the first-order term ends up as

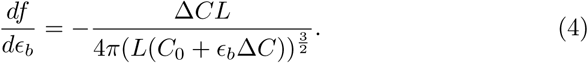

By using the values from testing, an inductance of 1 *µH*, a Δ*C* of around 5 *f F*, a *C*_0_ of 0.75 *pF*, and a relative bulk permittivity of 80, the sensitivity is approximately 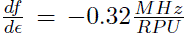 where *RPU* is the relative permittivity unit. The values for *C*_0_ and Δ*C* were determined by measuring the resonance frequency shift in deionized water, isopropyl alcohol (IPA), and in air using equation for an LC tank circuit, 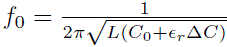. This approximation was verified by measuring the resonance frequency in water and IPA (a permittivity difference of around 60 RPU) and was found to determine the resonance frequency to within 5% of the measured value. Using the full RLC equation determined the resonance frequency even more precisely, as expected.

The limit of detection (LOD) of this device, defined here as 3*σ_n_* where *σ_n_* is the standard deviation of the resonance peak over time, was calculated to be 6.4 kHz, or 0.02 *RPU*. This calculation was based on the observation of the resonance shift over time of the sensor while immersed in the buffer solution. Drift was then corrected for and the standard deviation of the resonance frequency position was calculated. This gives the baseline noise from the sensing system, allowing us to find a minimum SNR of 3 times the noise standard deviation.

From the calibration curve shown in Fig. 4, the LOD is extrapolated to be around 7 nM for the tested antibody. This is calculated using the measured 3*σ_n_* of 6.4 kHz, and the calculated sensor sensitivity, 0.927 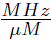.

Noise is the limiting factor in the LOD of electronic sensing systems and is typically driven by electrical interference or thermal effects. The majority of the noise in this system was observed to be from the VNA. It is possible that the sensor may also act as an antenna, picking up unwanted signals, and generating thermal noise, but this will mostly be mitigated by the scanning technique used by VNAs. The thermal noise generated by the sensor is also not anticipated to be significant since the sensor’s resistance is low at resonance and the bandwidth of the VNA is quite narrow when measuring S-parameters. In addition, it is possible to increase the input power when measuring the S-parameters, allowing for a higher SNR. This may appear as a simple fix, although it must be considered that the voltage at the nanogap under resonance conditions can be many times higher than the input voltage potentially causing the nanogap to become shorted and the sensor to be destroyed.

In addition to noise, signal drift can cause errors in measurement. Although this is generally easier to correct for in post-processing, it can still present significant issues in kinetic measurements. This is particularly an issue in FET-based sensors, as the threshold voltage is quite sensitive to temperature. The advantage of the STR architecture here is that, again, the sensing mechanism is passive. For example, there is no threshold drift that needs to be accounted for like in FET-based sensors. The primary concern in regards to drift at the sensor interface would be from thermal gradients slightly changing the permittivity of the solution under investigation. As for the VNA, this can be a concern as they are complicated devices. Particularly, the calibration of the VNA should be done same-day of the test being performed since aging the components can cause drift in the results. The device should also have sufficient time to warm up, as measurements may change over time due to the electronics approaching thermal equilibrium.

Since the BSA/anti-BSA system serves as a good model for proof of concept biomolecular kinetic measurements, many of these tests have been performed using optical methods. Various nanoplasmonic researcher groups have shown a typical LOD between 2.6 nM and 26 nM for IgG BSA antibodies [7, 45, 46].For example, Pang et al. created a two-dimensional nanohole array to serve as a sensor by passing the biofluid over the nanohole array. This sensor was tested using a BSA/anti-BSA system and revealed an LOD of 26 nM [7]. Additionally, Chung et al. investigated sensing multiple analytes using a single SPR chip by injecting different labels for the proteins under study. From the two methods they tested, the anti-BSA LODs were 21.3 nM and 2.6 nM [46]. The prototype STR architecture demonstrated here achieves an LOD of approximately 7 nM, a value certainly comparable to those reported from said research groups.

Although LOD is certainly an important metric, sensitivity must also be taken into account. To reasonably compare the sensitivities of STR and SPR, a normalized sensitivity to anti-BSA is reported in terms of *µM^−^*^1^. The system reported by Pang et al. is used here as they had a very similar solution under test and by extrapolating their values, the sensitivity to anti-BSA was determined to be around 3.68 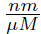 [7]. In order to compare directly with the STR architecture, the sensitivity needs to be normalized by the center wavelength of 1530 nm, giving approximately 0.0024 *µM^−^*^1^. Values extracted from plots in the study published by Chung et al. showed anti-BSA sensitivities to be around 7.5 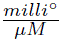 and 42 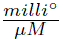 [46]. Normalizing these values by the center SPR angle of around 69.8*^◦^*, leads to normalized sensitivities of around 0.001 *µM^−^*^1^ and 0.006 *µM^−^*^1^. For the STR architecture, with a sensitivity of around 0.927 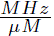 and a center frequency of around 150 MHz, the normalized sensitivity shows is calculated to be approximately 0.006 *µM^−^*^1^, again showing quite similar performance to these systems. This sensitivity metric, in addition to the LOD, shows that STR’s performance is on par with previous nanoplasmonic biosensors shown in research. Importantly, the presented STR prototype was engineered for portability and real-time phase detection, rather than maximizing sensitivity and LOD. With further optimization such as increasing input power, improving the Q-factor, narrowing the nanogap, and extending transmission line lengths, we anticipate further gains in performance. This lower LOD - in addition to the low-cost electronics, no moving optics, and a truly portable form factor - position STR as a competitive alternative for traditional gold standard SPR systems. Furthermore, as miniaturized RF electronics continue to advance, we envision future STR systems achieving further sensitivity gains through on-chip integration, improved Q-factors, and enhanced surface functionalization—potentially bringing SPR-class performance to every lab bench, clinic, and diagnostic kit.

In conclusion, we have demonstrated an STR biosensor that enables real-time, label-free molecular binding kinetics in a compact, palm-sized electronic platform. By leveraging nanogap-confined electric fields and operating at radio-frequencies, STR circumvents Debye screening, a typical obstacle for common electronic sensing methods. This architecture delivers quantitative kinetic measurements in physiologically relevant media - a capability traditionally reserved for optical gold standards such as SPR and fluorescence-based sensing.

Our results with BSA/anti-BSA specific interactions establish the viability of phase-resolved, frequency-based sensing as a robust, high-fidelity alternative to bulky optical systems. As Moore’s law continues to shrink electronics and enhance performance, STR offers a scalable path toward low-cost, portable, and integrable biosensors suitable for point-of-care diagnostics and lab-on-a-chip integration.

We envision future STR systems embedded in RF-ICs, enabling smart diagnostic wearables and field-deployable biosensing platforms. This work defines a new class of high-frequency, surface-specific electronic biosensor architecture, bringing SPR-like analytical power into the domain of microelectronics.

## Methods

### Sensor fabrication

The biosensor was fabricated on a microscope slide using photolithography, metal deposition, and focused ion beam (FIB). The substrate, a microscope slide, was baked at 150 C for 3 minutes. After, AZ1512 photoresist was spun on at 3000 rpm for 30 seconds. The resist was then exposed using a photolithography contact aligner. The sample was developed in AZ340:H2O (1:5) for 30 seconds before being washed. After, the sample was put into a reactive ion etching system where it was exposed to oxygen plasma with a flow rate of 99 sccm, a pressure of 100 mT, and a power of 100 W. The sample was put into plasma to improve the sidewall profile for cleaner liftoff. This sample was then put into a sputtering system where 10 nm of Ti and 175 nm of Au were deposited. Post metal deposition, the photoresist was lifted off using acetone.

To fabricate the nanogap, FIB milling was used. The gaps created were approximately 75 nm in width.

Although the sensors highlighted in this work were fabricated using FIB, which is not practical for large-scale manufacturing, there are methods, such as atomic layer deposition (ALD) lithography, that allow for devices of this type to be fabricated on large-scales [47–50]. ALD lithography utilizes the atomic resolution of ALD to create the minimum feature size, in this case the gap region. In conjunction with techniques for fabricating optical structures presented in [47–50], we developed a process to fabricate nanoscale gaps sufficient for sensing while being robust enough to support integration with electronics. The fabrication process, resulting SEM images, and *S*_11_ magnitude are shown in Fig. 5 and are able to be made at the wafer-scale, allowing for high-throughput and quick fabrication. Developing this wafer-scale fabrication process enables STR to integrate with RF-IC technologies and would further compact the sensing system. Future RF-IC technology integration may include a more optimized design to enhance sensor interfacing with the VNA. This would decrease device variability and increase device durability. This wafer-scale fabrication process is discussed in this paper’s Supplementary Material.

**Figure 5:**
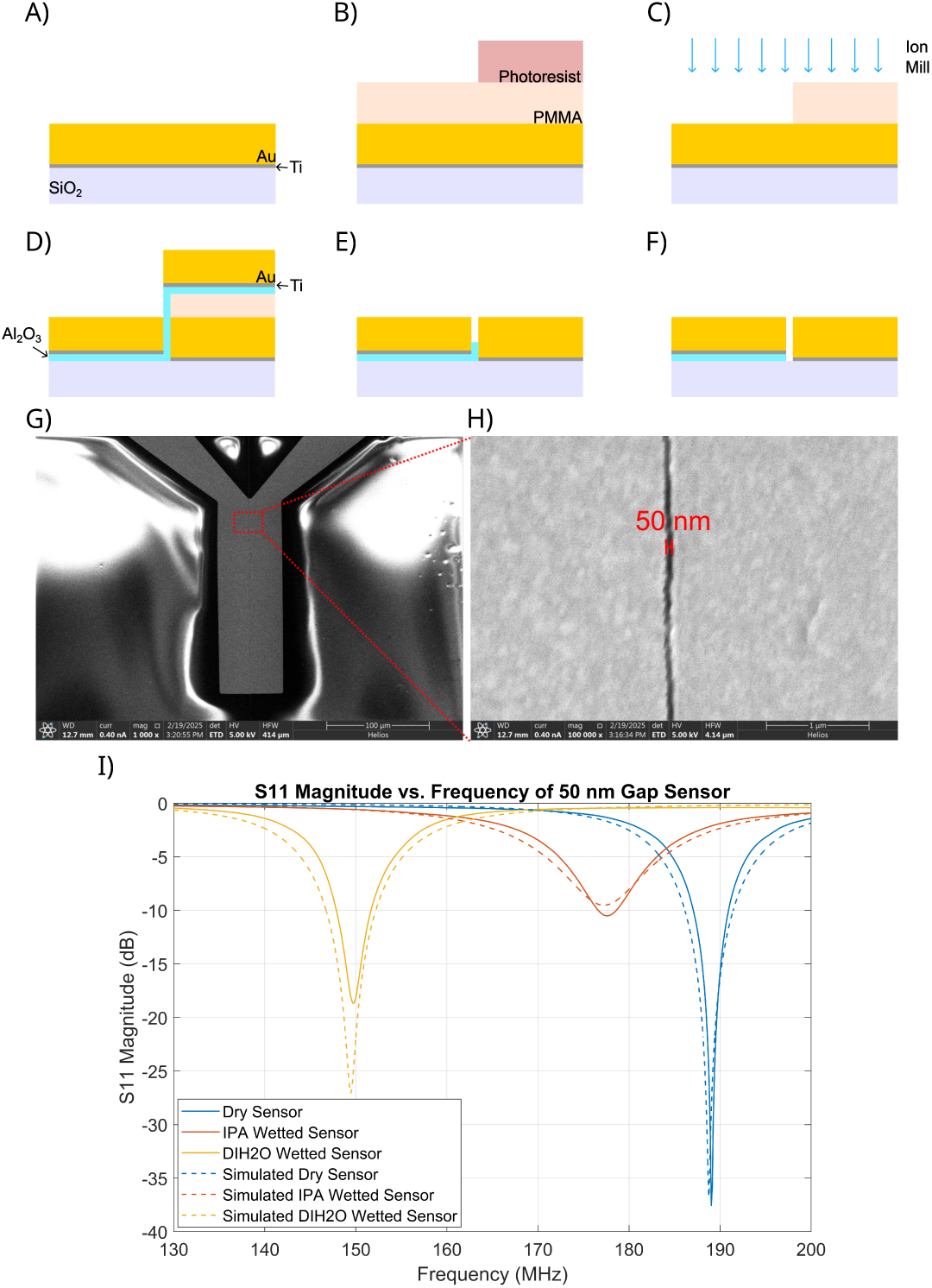
Wafer-scale fabrication process for developing nanoscale gaps for STR sensors and other various applications with ALD defining nanogap size. A) The substrate has been coated with Ti and Au, where Ti acts as an adhesion layer. B) PMMA and photoresist have been deposited on top of the metal and the photoresist was exposed with the desired pattern. The sample is exposed to oxygen plasma to etch away the PMMA and photoresist. C) The exposed metal is ion milled. D) Atomic layer deposition is used to define the gap width and, in this case, alumina is deposited. After ALD, Ti and Au are deposited and liftoff is performed. It was found during fabrication that most of the alumina within the gap was removed after liftoff, resulting in a functional sensor. E) The resulting structure can be exposed to an etchant to remove the alumina in the gap region if desired. F) The resulting structure and nanogap is formed. G) Resulting SEM of 200 *µ*m sensor region. H) SEM image of nanogap region. I) Measurement of *S*_11_ Magnitude of 50 nm gap sensor immersed in air, IPA, and water.

### Sensor assembly

After nanogap fabrication, PDMS microfluidic channels were fabricated and bonded to the sensors. Bonding required the samples, along with their respective microfluidic channels, to be placed in an oxygen plasma chamber with oxygen flowing at 30 sccm, and RF power of 150 W for 1 minute. After, the PDMS channels were firmly pressed down on top of the sensors, being careful that the channel was over the sensor’s electrodes and placed on a hot plate set at 75 C for at least 2 hours.

The final step included soldering on an inductor and SMA connector. These were soldered directly to the gold pads using solder containing 97% indium and 3% silver. Using an indium based solder prevents the gold from dissociating from the substrate too quickly, as more common tin-based solder quickly strip gold from the substrate [51]. Indium and gold still form an alloy, but this is a slower process.

### Data collection

The device used for conducting measurements was a nanoVNA V2 Plus4. This device was chosen, as it is affordable, handheld, and provides small step sizes at quick measurement rates. Resonance shifts were determined from the measured *S*_11_ magnitude and phase after calibration. The open source software NanoVNA-QT was used to conduct measurements and save S-parameter data and a MATLAB script was created to read and process the data.

To test whether protein binding kinetics could be measured, bovine serum albumin (BSA) was dissolved in a 0.1x phosphate buffered saline (PBS) solution at a concentration of 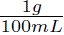 to form a 1% solution. This was injected into the microfluidic channel to form a BSA monolayer over the sensor’s surface. After the BSA blocking step was completed, a solution containing Anti-BSA IgG antibody (purchased from Thermo Fisher Scientific Product # A11133) was diluted to one tenth of its original concentration with deionized water to achieve a PBS concentration of 0.1x. For further dilution steps, the 0.1x PBS solution created in the lab was utilized. After dilution, the solution was then injected and the response was observed. This whole process is shown in Fig. 2 and the resulting resonance shift over time is shown in Fig. 4.

This process was repeated for multiple antibody concentrations to determine association and dissociation constants and was verified using SPR.

In order to determine the sensitivity of each sensor, IPA was injected into the sensor and the resonance shift was recorded. This was done in addition to injecting deionized water. The permittivity of these substances is readily known, so the shift in the resonance frequency allowed for *C*_0_ and Δ*C* to be calculated. This step allowed for a more accurate concentration of antibodies to be determined and to verify the underlying principles of the sensor’s operation, but is not necessary for sensing applications which are not quantitative.

## Data availability

The datasets used and/or analyzed during the current study are available from the corresponding author upon reasonable request.

## Code availability

The underlying code for this study is available at: https://github.com/bry1275/STR_data_analysis. An example dataset is provided.

## Acknowledgments

This research was supported by the U.S. National Science Foundation (Grant No. ECCS 2240245). S.-H.O. further acknowledges support from the Sanford P. Bordeau chair.

## Author contributions

S.-H.O. and B.K.C. conceived and designed the study. B.K.C. designed, fabricated, and tested the sensors. S.-H.O. and B.K.C. wrote the manuscript. B.K.C. visualized the data and designs. S.-H.O. and B.K.C. discussed the results. All authors reviewed the manuscript. All authors approved the manuscript.

