## Supplementary material for "Surface Transmon Resonance (STR): a handheld nanogap biosensor for real-time, label-free molecular binding kinetics"

#### Sensor operating principles

When the solution is conductive, a conductance term, $G$ in units of siemens, must be taken into account for calculating the resonance frequency. The resonance frequency for a conductive medium is then

$$f_0 = \frac{1}{2\pi}\sqrt{\frac{1}{LC} - \frac{G^2}{C^2}}. \tag{1}$$

Since the gap between transmission lines is so small, the electric field is confined to the nanogap and the capacitance can be modeled as a parallel-plate capacitor $\Delta C = \frac{\epsilon_0 A}{g}$ , where $\Delta C$ is the capacitance of the nanogap when empty, $\epsilon_0$ is the vacuum permittivity, $A$ is the surface area of the electrodes, and $g$ is the distance between the electrodes. The change in this capacitance term is what primarily determines the shift of the resonance peak, which, in turn, is mainly driven by the relative permittivity of the medium within the gap region. The permittivity of the gap region is then determined by the volumetric fraction of biomolecules within the gap by the equation

$$\epsilon_r = \epsilon_b + \frac{N_m V_m}{V_g}(\epsilon_m - \epsilon_b). \tag{2}$$

$\epsilon_r$ is the relative permittivity within the gap, $\epsilon_b$ is the bulk permittivity or permittivity of the fluid, $N_m$ is the number of molecules within the gap region, $V_m$ is the volume of a single molecule, $V_g$ is the volume of the gap region, and $\epsilon_m$ is the permittivity of a single molecule at the measurement frequency.

The equation can be rearranged to provide an estimate of the number of biomolecules within the gap region. This resulting equation would be

$$N_m = \frac{V_g}{V_m}\frac{\epsilon_r - \epsilon_b}{\epsilon_m - \epsilon_b}. \tag{3}$$

If these equations are used analytically, it is important to note that they do not consider the conductance change of solutions and may be inaccurate if this isn't taken into account.

For phase-based measurements, the resonance frequency is determined from where the $S_{11}$ phase crosses zero. This zero-point can be determined by interpolation of the $S_{11}$ data, or calculation based on equation (10). The $S_{11}$ parameter is calculated from the characteristic impedance of the Vector Network Analyzer (VNA) and the input impedance of the sensor system, shown in equation (5), and is typically a complex value. To determine the $S_{11}$ parameters of the sensor, the impedance of a series RLC circuit is used to model the sensor and defined as

$$Z_{in}(\omega) = R + j(\omega L - \frac{1}{\omega C}), \tag{4}$$

where $R$ is the series resistance, $j$ is the imaginary number, $\omega$ is the angular frequency, $L$ is the inductance, and $C$ is the capacitance of the sensor. $S_{11}$ is measured as a reflection coefficient in this case, as there is only one port, and is determined by the VNA's characteristic impedance by

$$S_{11}(\omega) = \frac{Z_{in}(\omega) - Z_0}{Z_{in}(\omega) + Z_0}. \tag{5}$$

In order to determine a resonance peak accurately, a sharp resonance peak and steep phase change is desired. It isn't immediately obvious from these equations but, by combining them and taking the first few terms of the Taylor expansion in terms of $Z_{im}$, the optimal operating point is more clear. Since the first derivative of $|S_{11}|$ is equal to 0 near resonance, the second derivative is taken. We are also mainly concerned about what happens at resonance, so $Z_{im}$ is set to 0 in the zeroth and higher order terms.

To determine the Taylor expansion of $S_{11}$'s magnitude and phase, equations (4) and (5) are combined to give

$$S_{11} = \frac{Z_{im}^2 + Z_r^2 - Z_0^2 + j2Z_0Z_{im}}{(Z_r + Z_0)^2 + Z_{im}^2}, \tag{6}$$

where $Z_{im}$ is the imaginary part of the impedance and $Z_r$ is the real part of the impedance. This equation can be then separated into the magnitude and phase, providing a direct comparison to measured values

$$|S_{11}| = \frac{\sqrt{Z_r^4 + Z_0^4 + Z_{im}^4 - 2Z_0^2Z_r^2 + 2Z_r^2Z_{im}^2 + 2Z_0^2Z_{im}^2}}{(Z_r + Z_0)^2 + Z_{im}^2}, \tag{7}$$

$$\phi_{11} = \arctan\left(\frac{2Z_{im}Z_0}{Z_r^2 - Z_0^2 + Z_{im}^2}\right). \tag{8}$$

The first derivative of the phase as well as the second derivative of the magnitude can be taken to determine the Taylor expansion. The first derivative of the magnitude ends up as 0, so that term is neglected. The Taylor series ends up as

$$|S_{11}| \approx \begin{cases} \dfrac{Z_r - Z_0}{Z_r + Z_0} - \dfrac{2Z_0 Z_r}{(Z_r + Z_0)^2 |Z_r^2 - Z_0^2|} Z_{im}(\omega)^2 & Z_r < Z_0, \\ \dfrac{Z_r - Z_0}{Z_r + Z_0} + \dfrac{2Z_0 Z_r}{(Z_r + Z_0)^2 |Z_r^2 - Z_0^2|} Z_{im}(\omega)^2 & Z_r > Z_0, \end{cases} \tag{9}$$

for the magnitude of $S_{11}$ and

$$\phi_{11} \approx \begin{cases} -\pi + \dfrac{2Z_0}{Z_r^2 - Z_0^2} Z_{im}(\omega) & Z_r < Z_0, \\ \dfrac{2Z_0}{Z_r^2 - Z_0^2} Z_{im}(\omega) & Z_r > Z_0, \end{cases} \tag{10}$$

for the angle of $S_{11}$ in radians.

With these approximations, it is now quite clear that as $Z_r$ approaches $Z_0$, the resonance peak sharpens and the phase becomes steeper, thus allowing for the increase of accuracy in the measurements. It also becomes obvious that a conductive solutions will directly change the magnitude's sharpness and phase's steepness because $Z_r$ will change.

Determining $C_0$ requires more complex transmission line analysis. A reasonable portion of the stray capacitance is from the transmission lines leading from the SMA connector to the sensing region. This capacitance can be calculated using transmission line equations for coplanar electrodes. First, the equation for the relative permittivity surrounding the transmission line is

$$\epsilon_r' = \frac{\epsilon_r + 1}{2} \{\tanh(0.775 \ln(\frac{h}{w}) + 1.75) + \frac{kw}{h}(0.04 - 0.7k + 0.01(1 - 0.1\epsilon_r)(0.25 + k))\}. \tag{11}$$

In this equation, $h$ is the height of the substrate, $w$ is the width of the transmission lines, and $\epsilon_r$ is the permittivity of the substrate. The variable $k$ follows the equation

$$k = \frac{s}{s + 2w}, \tag{12}$$

where $s$ is the distance between the transmission lines. From here, the capacitance per unit length is calculated by

$$c' = \begin{cases} \dfrac{\epsilon_r'}{120\nu_0 \ln(2\frac{1+\sqrt{k}}{1-\sqrt{k}})} & \dfrac{1}{\sqrt{2}} \leq k \leq 1, \\ \dfrac{\epsilon_r' \ln(2\frac{1+\sqrt{k'}}{1-\sqrt{k'}})}{377\pi\nu_0} & 0 \leq k \leq \dfrac{1}{\sqrt{2}}, \end{cases} \tag{13}$$

where $\nu_0$ is the speed of light, and $k'$ is defined by $k' = \sqrt{1 - k^2}$. The resulting capacitance can then be multiplied by the length of the lines to get the stray capacitance contributed by the transmission lines. From measured data, this values ends up around 3x less than the actual value of the stray capacitance. This is most likely due to parasitic capacitances from solder joints and non-ideal parts such as the SMA connector and inductor.

#### Wafer-Scale nanogap electrode fabrication process

For the nanogap fabrication, the substrate starts off by undergoing a dehydration bake at 200 C for 10 minutes before depositing metal. For the first metal stack, 10 nm Ti is deposited and 200 nm of Au. The specific thickness of Au along with the ALD film will determine the effective sensitivity of the resulting STR sensor. After metal deposition, the sample is then coated with 950 PMMA C9 and spun at 4000 rpm for 1 minute. It is then baked again at 200 C for 8 minutes to drive out any residual solvents. Afterwards, the sample undergoes the photolithography process previously described for the FIB samples. To transfer the photoresist pattern into the PMMA, the sample is loaded into an oxygen plasma chamber and exposed to oxygen plasma with a flow rate of 99 sccm, a pressure of 100 mT, and an RF power of 100 W. The bilayer process of the PMMA and photoresist is required since loading the photoresist into the thermal ALD chamber would cause contamination to the system. PMMA is able to be baked at a higher temperature which doesn't cause contamination of the thermal ALD chamber since the solvents are driven out in the 200 C bake step. Once the pattern is transferred to the PMMA layer, the sample can be ion milled. After the ion mill etches through the titanium adhesive layer, the sample can be loaded into an ALD chamber. The ALD system utilized in this paper was the Ultratech Savannah-Thermal ALD system, and alumina was deposited at 180 C. The number of cycles completed in the ALD system determines the resulting gap width. After ALD, the second metal layer can be deposited. Our technique uses the 10 nm Ti adhesion layer to maintain robustness when soldering. The gold thickness after this step can be determined by subtracting the first layer's gold thickness by the ALD film's thickness. Once metal deposition is complete, liftoff can be performed. Our process showed best results when liftoff was performed in acetone for around 1 to 2 days. It seemed, after liftoff, that the majority of the alumina with the gap had also been removed. A cross section of this is shown in Supplementary Fig. S1. To get rid of any excess alumina in the gap, wet etching with an H3PO4-based etchant.

For further definition of the nanogap structure, the photolithography process can be repeated with the desired pattern and the sample can be placed in the ion mill. If there are electrical shorts present, a glancing angle ion milling process can greatly improve electrical isolation. It is important to note that glancing angle ion milling will still vertically etch the gold, although at a much slower rate. After glancing angle ion milling, there were occasionally some sensor still shorted. It was tested and found that, by applying a low voltage to the electrodes, around 1-2 V, the small shorts end up vaporizing and isolating

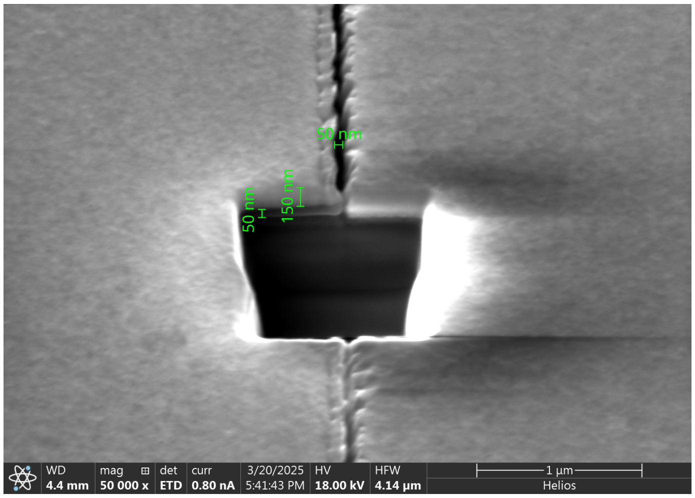


Figure S1: SEM image of gap made from wafer-scale fabrication process post liftoff. The gap was cut using FIB in order to see the cross sectional-profile. From this perspective, the partial removal of the alumina, due to liftoff, is visible. Scale bars are added from SEM/FIB measurements. The 50 nm alumina layer can be seen underneath the 150 nm Au/Ti electrode. The gap also has the same width as the ALD layer thickness of 50 nm.

each electrode. A small voltage is used since higher voltages can destroy the electrodes. The smallest gap size attempted with this process was 25 nm and the resulting STR sensor was operational.

### SPR verification of protein kinetics

SPR was used to verify the binding coefficients of the BSA antibody. BSA was deposited on the surface, similar to the STR experiments, and 0.002% anti-BSA was used to determine the protein kinetics. Typically this is done with multiple concentrations, but there was only enough antibody from the batch to conduct this measurement. Despite only having one SPR curve, very reasonable values were determined with very good fits ($R^2 > 0.995$). From this curve, $K_D$ was determined to lie between 100 - 160 nM given the 95% confidence bounds from the fits.

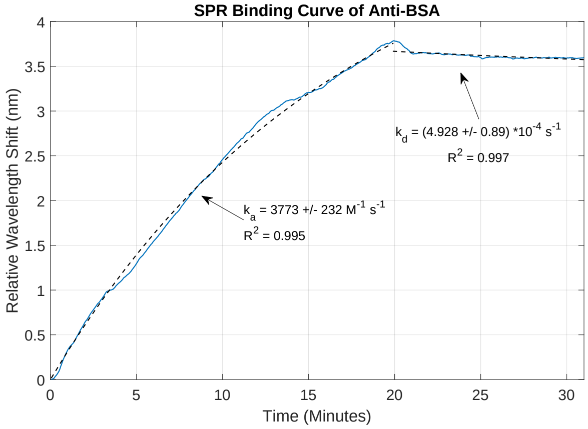


Figure S2: SPR curve from 0.002% anti-BSA concentration showing binding and unbinding coefficients with 95% confidence interval.
